# Convergent Functional Constraints Make Biochemical Reaction Diversity Predictable

**DOI:** 10.1101/2025.06.20.660822

**Authors:** Veronica V. Mierzejewski, Cole Mathis, Christopher P. Kempes, Sara I. Walker

## Abstract

The predictability of biochemical evolution remains highly debated, as it is unknown if biological function can be predicted *a priori*. Here, we explore predictability of function not in individual reactions, but in ensembles. Sampling from 13,777 bacterial, 371 archaeal, and 203 eukaryotic genomes, we use Enzyme Commission (EC) classification to hierarchically group enzyme-catalyzed reactions that perform similar transformations into functional equivalence classes. We show that organisms partition their reaction diversity among these functional equivalence classes in a predictable way, identifying over 120 new, system-size dependent functional scaling relationships. We find distantly related lineages, acting on distinct molecular substrates, can display similar functional scaling, demonstrating that convergence is not driven by phylogeny or common reactions. We demonstrate how transitions in functional scaling can be identified with physiological shifts, using as case studies O_2_-utilizing oxidoreductases and hexosyltransferases. Taken together, our results open novel avenues for predicting global features of evolving enzyme populations, independent of protein structures. These functional constraints may have broad implications, including for predicting biochemical diversity, designing synthetic organisms, and modeling the evolution of reaction diversity in cases where the exact identity of catalysts is not known, such as during the emergence of life and for potential alternative forms of biochemistry.

## INTRODUCTION

The predictability of evolution is highly debated, especially whether it is possible to predict biological functions *a priori*^1–3^. Well-known examples of technical and fundamental limits in predictability can be found in enzyme structure and function. Enzymes, which catalyze most biochemical reactions, evolve along historically contingent paths of mutations and recombinations. While these changes occur at the sequence level, innovation in function arises from protein structure. Within the cell, a single protein sequence manifests as a narrow ensemble of structural conformations. Predicting distributions of these conformations over successive mutation events is not possible: statistical thermodynamics places hard physical limits on predicting how these distributions will change along molecular lineages^4^, meaning protein structure, and therefore function, cannot be predicted. Furthermore, the genetic background of the cell will determine whether a mutation is advantageous, deleterious or neutral, an effect known as epistasis^5,6^. The interplay between the evolution of sequence, and selection on structure, thus leads to trajectories of macromolecular evolution that are generally unpredictable at the scale of individual protein catalytic function.

Despite these limitations, there are cases where molecular function is so strongly convergent as to suggest the possibility for prediction. A prominent example is the evolution of C4 photosynthesis as an adaptive response to dry climates. Evolving independently >45 times via different chemical and anatomical means^7^, C4 photosynthesis demonstrates how living systems can decouple functions from precise molecular instantiations to achieve convergence^8^. Indeed, C4 photosynthesis provides an explicit example of *multiple realizability*, where different molecules or mechanisms that are *functionally equivalent* (i.e., sharing the same roles) can lead to the same outcome^9,10^. Functional equivalence classes are sets of molecules and reactions with the same function. Identifying functional equivalence classes, and what factors shape them, could enable powerful predictive tools in cases where it is not possible to predict evolution in individual molecules. Thus, even if it is not possible to forecast precisely which biochemical reactions evolution will discover, it may still be possible to use functional constraints to illuminate the space of possible solutions^11^.

Enzymes provide an opportunity to test this hypothesis. Enzyme Commission (EC) numbers are four-digit codes (EC -.-.-.-) that group enzyme-catalyzed reactions hierarchically into categories based on reaction chemistry, where each successive digit provides a progressively finer classification of the specified reaction^12^. The 1^st^ digit groups enzymes into six broad reaction classes: redox (**EC 1**), transferase (**EC 2**), hydrolase (**EC 3**), lyase (**EC 4**), isomerase (**EC 5**) and ligase (**EC 6**), which cover the major categories of catalyzed reactions found in living systems. We do not include the newly established category of translocases (EC 7), as these functions can be described by the other reaction classes (e.g., EC 7.1.1 are oxidoreductase reactions). Intermediate levels in the hierarchy (2^nd^ and 3^rd^ digit) only identify a few substrates, while a full four-digit EC code specifies all reactants and products of a given reaction. A four-digit EC code uniquely specifies a reaction, but importantly, it does not uniquely specify an enzyme catalyst, as different enzymes with distinct sequences and structures can evolve to catalyze the same reaction. In this way, *every* level of the EC hierarchy can represent a functional equivalence class, corresponding to many possible protein catalysts. For example, EC 2.1.1.5, a transferase reaction that transfers a methyl group from betaine to L-homocysteine to form the amino acid L-methionine, represents a functional equivalence class because at least two different enzymes can catalyze this reaction^13^. Moving up the EC hierarchy, we can group reactions into broader functional equivalence classes that share common transformations but become increasingly abstracted from substate-specificity: continuing on our example, **EC 2.1.1.-**are a set of 376 reactions that transfer methyl groups between any pair of molecules; the class **EC 2.1.-**consist of 407 reactions that transfer any one-carbon group between any pair of molecules; and **EC 2** represents the 2085 total biochemical transferase reactions cataloged to date^14^.

Biochemical reactions partitioned into broad classes at the 1^st^ EC digit exhibit system-size dependent scaling trends, which were recently shown to be universal across archaeal, bacterial and eukaryotic lineages^15^. However, it is not clear why there should be such consistent scaling constraints on global reaction diversity, especially considering (1) there is a vast diversity of catalytic protein structural domains cataloged to date^16–18^, 2) there are almost no substrate-level constraints on reactants or products for 1^st^ digit ECs, and (3) reactions within the same 1^st^ digit EC can play completely different biological roles in the cell. One hypothesis is that scaling trends reflect shared ancestry, ultimately rooting all biochemical similarity at the last universal common ancestor (LUCA). Alternatively, these scaling laws may provide evidence of convergent evolution in biochemistry, where patterns in reaction diversity are strongly convergent due to environmental or physiological constraints. If this alternative hypothesis is true, then such scaling relationships could provide new predictive tools for the possibility space of reactions that can be selected in evolution.

We use the functional equivalence grouping of enzymes across the EC hierarchy to probe the role of homology versus convergence in reaction diversity. To this end, we identify 120 new scaling trends to quantify similarities and differences in the use of functionally equivalent sets of reactions across taxa at different degrees of taxonomy and reaction specificity. We identify differences in functional scaling across the EC hierarchy and show these are not driven by phylogeny, nor by the presence of shared reactions across organisms. We show transitions in functional scaling can be identified with physiological shifts. Our results support the conclusion that cellular context, including physiology and potentially environmental factors, constrain the reaction diversity evolutionarily accessible to viable organisms. This leads to the surprising consequence that functional compositions of evolving reaction networks may be predictable in their ensemble properties, with implications for designing catalyzed systems in bioengineering^19,20^, or predicting evolving systems where the exact identity of catalysts might be unknown, including in emergence of life research^21,22^ and in astrobiology^23^.

## RESULTS

### Functional Scaling Arises from Convergence, Not Shared Ancestry

We track the diversity of reactions for functionally annotated enzymes, represented as four-digit EC codes, by sampling 13,777 bacterial, 371 archaeal, and 203 eukaryotic genomes from the Joint Genome Institute’s Integrated Microbial Genomes and Microbiomes (DOE-JGI IMG/M) database^24,25^. We group reactions into hierarchical groups designated by 1^st^, 2^nd^, and 3^rd^ digit EC levels to define *functional equivalence classes* of varying degrees of substrate-specificity. We make one modification to the EC nomenclature and group together **EC 3** (hydrolases) and **EC 4** (lyases), which are only differentiated by the presence of a single substrate: H_2_O, **Table 1**. Because our aim is to identify substrate-agnostic regularities based on equivalent function, we combine these two sets of reactions and label the functional equivalence class of **EC 3/4** as ‘lyases’ throughout the text. To track reaction diversity in a given functional equivalence class, we bin reaction data by counting all reactions in an organism up to a specified digit (e.g., counting all four-digit EC codes that start with ‘1’ to count all oxidoreductases, all four-digit EC codes that start with ‘1.2’ to count all oxidoreductases that act on the aldehyde or oxo group of electron donors, etc.). Scaling behavior for a functional equivalence class is then identified by determining how the number of unique reactions in each grouping change with system size, which we designate as *functional scaling*. Here, ‘size’ is the total number of unique reactions across all EC codes identified in a genome. Functional scaling elucidates whether the diversity within a related group of reactions grows disproportionately larger (superlinear, with scaling exponent >1), disproportionately smaller (sublinear, with scaling exponents <1), or at constant proportion (linear, where scaling exponents are statistically indistinguishable from 1) as the total number of reactions encoded in genomes increases.

**Table 1:**
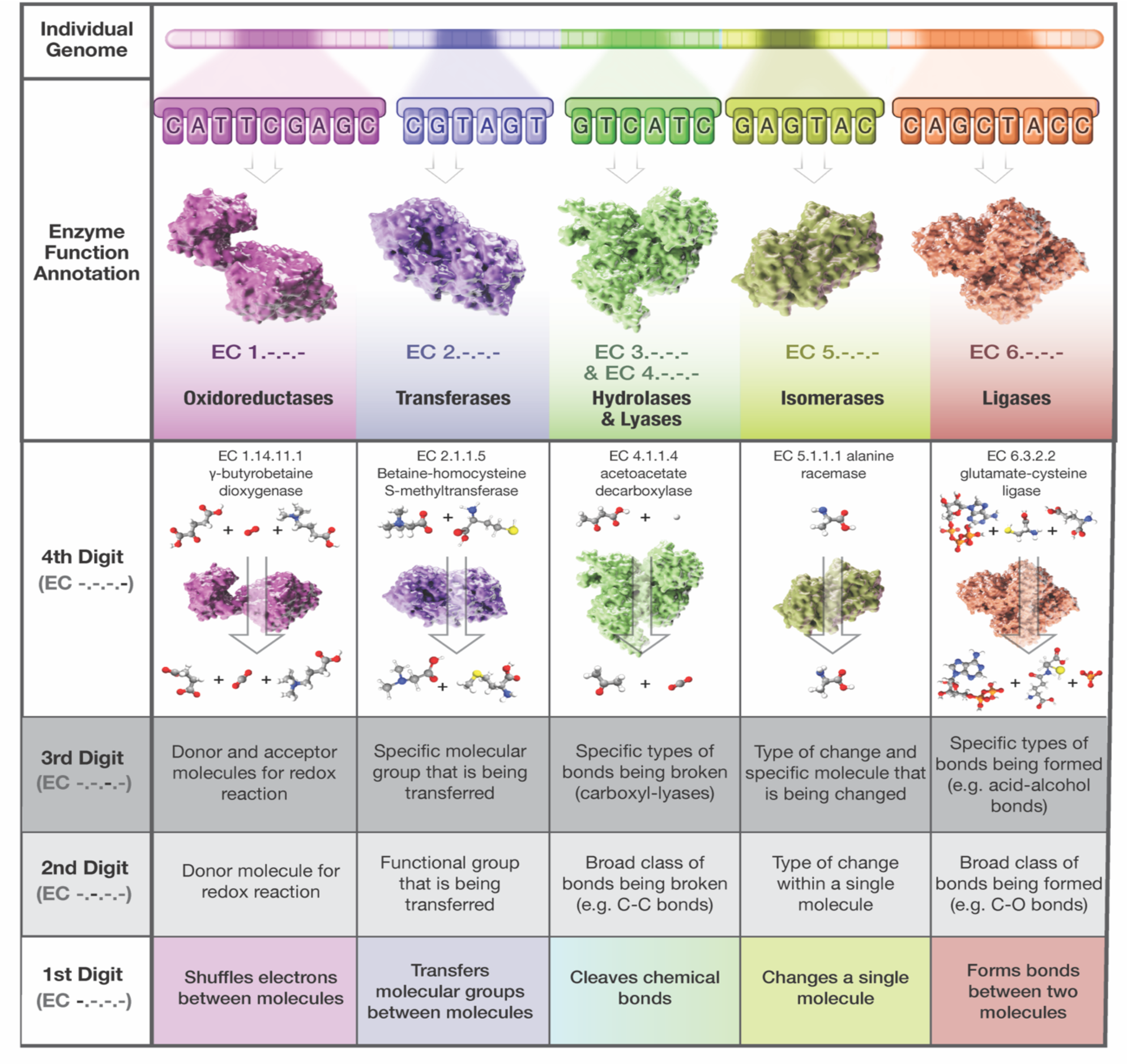
The Enzyme Commission (EC) hierarchy can be used to study functional equivalence classes of enzyme catalyzed reactions. Functionally annotated genomes from the Joint Genome Institute’s Integrated Microbial Genomes and Microbiomes (DOE-JGI IMG/M) database allow for identifying four-digit EC codes (EC -.-.-.-) that uniquely specify genomically encoded reactions. Reactions identified within each taxon are placed in hierarchical groups designated by 3^rd^, 2^nd^ and 1^st^ digit EC groupings to define *functional equivalence classes* of varying degrees of substrate-specificity. We group together 1^st^ digit EC codes 3.- and 4.-, corresponding to hydrolases and lyases, respectively, because these two broad reaction classes are only differentiated by the presence of one substrate, water. 1^st^, 2^nd^ and 3^rd^ digit groupings only identify a few substrates, while a full four-digit EC code specifies all reactants and products of a given reaction. Thus, a four-digit code represents a functional equivalence class of enzyme catalysts, whereas fewer digits identify functional equivalence among reaction substrates.

To test hypotheses about the role of homology versus convergence in reaction diversity, we partition our data in two ways. The first is phylogenetically, **Figure 1A**, in which we calculate scaling parameters across different phyla. Studying functional scaling relationships at this level allows us to determine how system size-dependent patterns in reaction use may or may not be driven by evolutionary proximity. The second way is by level of the EC hierarchy, **Figure 1B**, to explore a larger set of functional scaling relationships from which underlying physiological explanations can start to be mapped.

**Figure 1:**
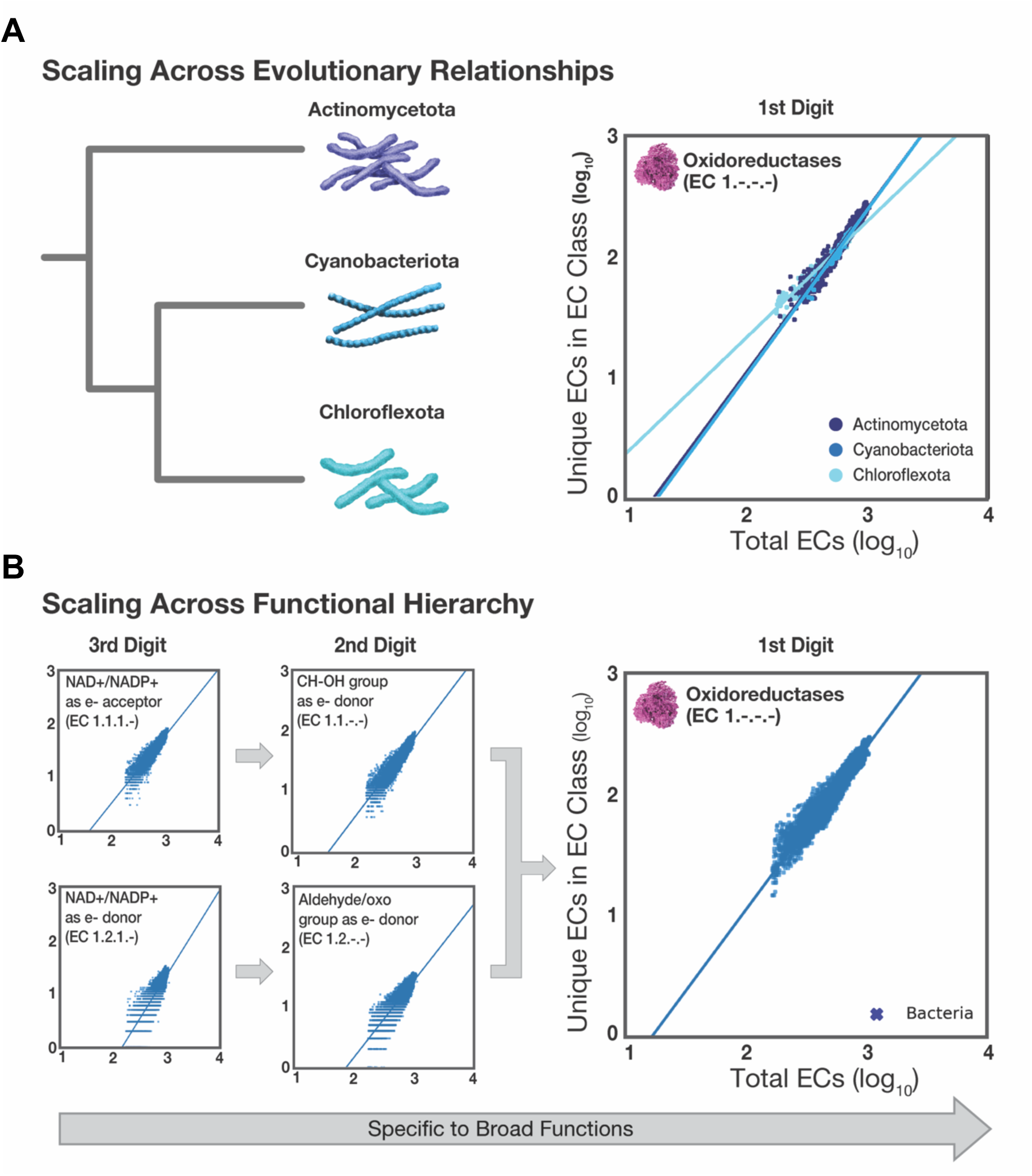
Predicting biochemical reaction diversity with scaling relationships that describe membership in functional equivalence classes. (A) Similarity in functional scaling may be the result of phylogeny. We test hypotheses of evolutionary homology versus convergent evolution by correlating scaling coefficients with evolutionary proximity (empirical data used). (B) Functional scaling laws can be identified at varying levels of the EC hierarchy. If the presence of some of these functional scaling relationships are shown to be convergent and not dependent on shared phylogeny nor reaction similarity, exploring functional scaling relationships at a finer-grained level might reveal physiological drivers of convergent behaviors.

We first examine functional scaling trends to determine if they are driven by homology or arise through evolutionary convergence. To do so, we first group enzymes by the 1^st^ EC digit across 16 bacterial phyla.

We identify 85 new functional scaling trends at the phylum-level. We find these scaling exponents are mostly consistent with domain-level scaling trends^15^ (**Figure 2A, SI Data Files S1 and S2**); however, deviations from this nearly universal behavior are also apparent. Transferases (**EC 2**) exhibit the most fidelity to their domain-level scaling, indicating that the diversity of transferase reactions is much more constrained than the diversity of other functional equivalence classes, such as ligases (**EC 6**), which vary more than the other classes. We performed permutation tests (**SI Figure S1**), confirming that variation in phylum-level functional scaling trends is statistically significant and not explained solely by domain-level data.

**Figure 2:**
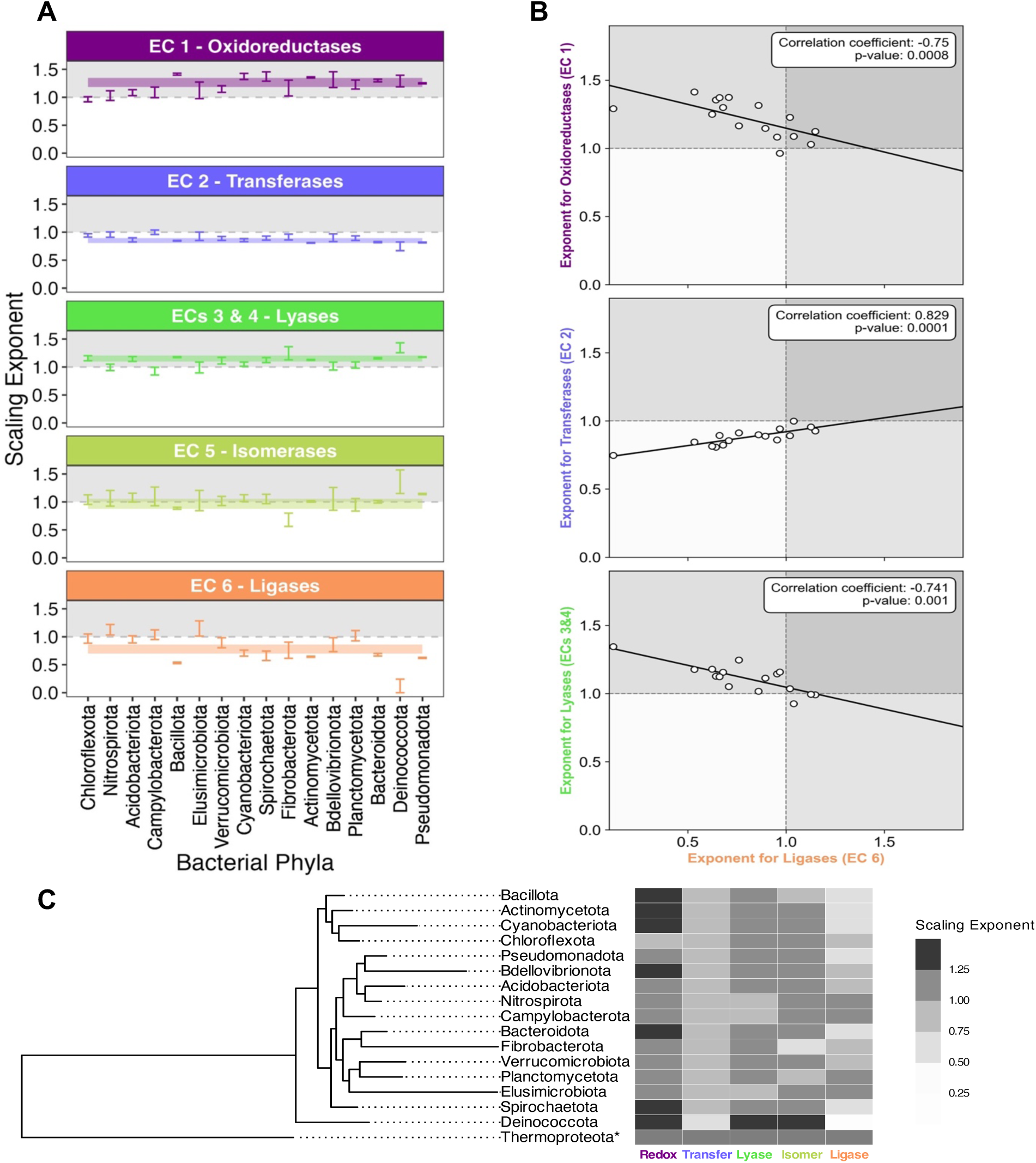
Functional scaling reflects constraints on reaction diversity that are not driven by shared ancestry. **(A)** Scaling exponents for bacterial phyla calculated for each 1^st^ digit EC functional equivalence class. Error bars indicate 95% confidence intervals. **(B)** Correlations among scaling exponents for oxidoreductases, transferases, lyases, and ligases across phyla, show system size-dependent trade-offs in allocation to different classes. **(C)** An adapted tree from Martinez-Gutierrez and Aylward, with corresponding scaling exponents for each class, show a lack of correlation between scaling and evolutionary relationships.

Because functional interdependencies are ubiquitous in cells^26–28^, we asked if variation in functional scaling might reveal previously uncharacterized interdependencies in reaction diversity across EC 1^st^ digit classes. To determine this, we examined if scaling exponents for the different ECs are correlated. We find that in several cases, there are statically significant correlations, with ligases (**EC 6**) implicated in all of them (**Figure 2B)**. The scaling exponents of oxidoreductases (**EC 1)** and lyases (**EC 3/4**) are negatively correlated with those of ligases, indicating how phyla whose genomes tend to encode more oxidoreductase or lyase reactions with size do so at the expense of encoding fewer ligase reactions. The negative correlation between lyases, which split molecules, and ligases, which combine them, is particularly surprising. *A priori*, one might expect this functional opposition to lead to trends where these two functions increase in tandem in a homeostatic fashion, but this is not what we observe. It is possible that this and other identified trade-offs may reflect ecological niche (e.g., some phyla being predominantly consumers that break down biomass, requiring more lyase reactions). We find one functional equivalence class, the transferases (**EC 2**), are positively correlated with ligases, which may be due to their cooperation in constructing larger molecules. Notably, although scaling exponents for oxidoreductases, lyases, and transferases are all correlated to that of ligases, these classes are not correlated with each other (**SI Figure S2**), implying that the above correlations are not simply a result of redistributing a fixed number of reactions to different functional classes, but instead reflect ensemble constraints tied to ligase diversity.

We next analyzed scaling exponents across phyla with respect to their position in an adapted tree of life, derived from Martinez-Gutierrez and Aylward^29^ (**Figure 2C)**. Our tree shows phylum-level relationships, and we include Thermoproteota, an archaeal outgroup to probe trends across large evolutionary distances. Overall, we see no correlation between phylogenetic distance and scaling exponent across any 1^st^ digit ECs, as confirmed by p-values below 0.01 in Spearman correlation tests (**SI Figure S3, SI Table S1**). We do see cases where functional scaling can vary substantially across the tree. For instance, the sublinear scaling observed for oxidoreductases in Chloroflexota differs dramatically from the superlinear patterns in its closest evolutionary neighbors, the Bacillota, Actinomycetota, and Cyanobacteriota, as well as in bacteria overall. Additionally, we observe distant phyla have converged to similar scaling exponents: Nitrospirota, Campylobacterota, Planctomycetota and Elusimicrobiota all display superlinear scaling in ligase reactions, despite their distance from one another on the tree and their closer evolutionary neighbors displaying sublinear scaling. This is also true across biological domains: functional scaling for our outgroup, the Thermoproteota archaea, is similar to that of many bacterial phyla in one or more functional class. We therefore conclude that the factors shaping functional scaling are not driven by shared ancestry.

### Functional Scaling is Not Dictated by Substrate

We ask if the observed convergent functional scaling trends represent truly emergent patterns**—**where different sets of reactions lead to the same functional scaling**—**or if they instead reflect organisms relying on the same reactions (**Figure 3A)**. In other words, we ask: are constraints in reaction diversity multiply realizable? To explore the possibility of multiple realizability of functional scaling, we quantified the average reaction set similarity between taxa by identifying how many four-digit EC codes they share (see **Materials and Methods**). We find similarity in functional scaling is not fully determined by shared reactions across taxa (**Figure 3B)**. Supporting multiple realizability, taxa can be >50% different from one another in terms of encoded reactions, while displaying similar functional scaling behavior, which we see examples of for most broad functional equivalence classes. Oxidoreductases and lyases show the largest differences, with oxidoreductase reaction sets having average similarities as low as 30% and lyase sets as low as 40%, while ligases show the highest similarities (above 80%, even between evolutionarily distant phyla). Intriguingly, the low similarities between bacterial and archaeal phyla are not a consequence of which reactions are available to each domain: nearly all reactions found in archaea are also in bacteria (**Figure 3C)**. This similarity could reflect the under-sampling of archaea relative to bacteria^30^, but also indicates that despite similarity in reactions at the domain-level, the reaction sets of individual phyla between archaea and bacteria, representing the composition of viable reaction networks, can differ substantially. These data suggest convergence in functional scaling relationships is not dictated by specific reactions but instead reflects emergent constraints at the level of ensemble diversity.

**Figure 3:**
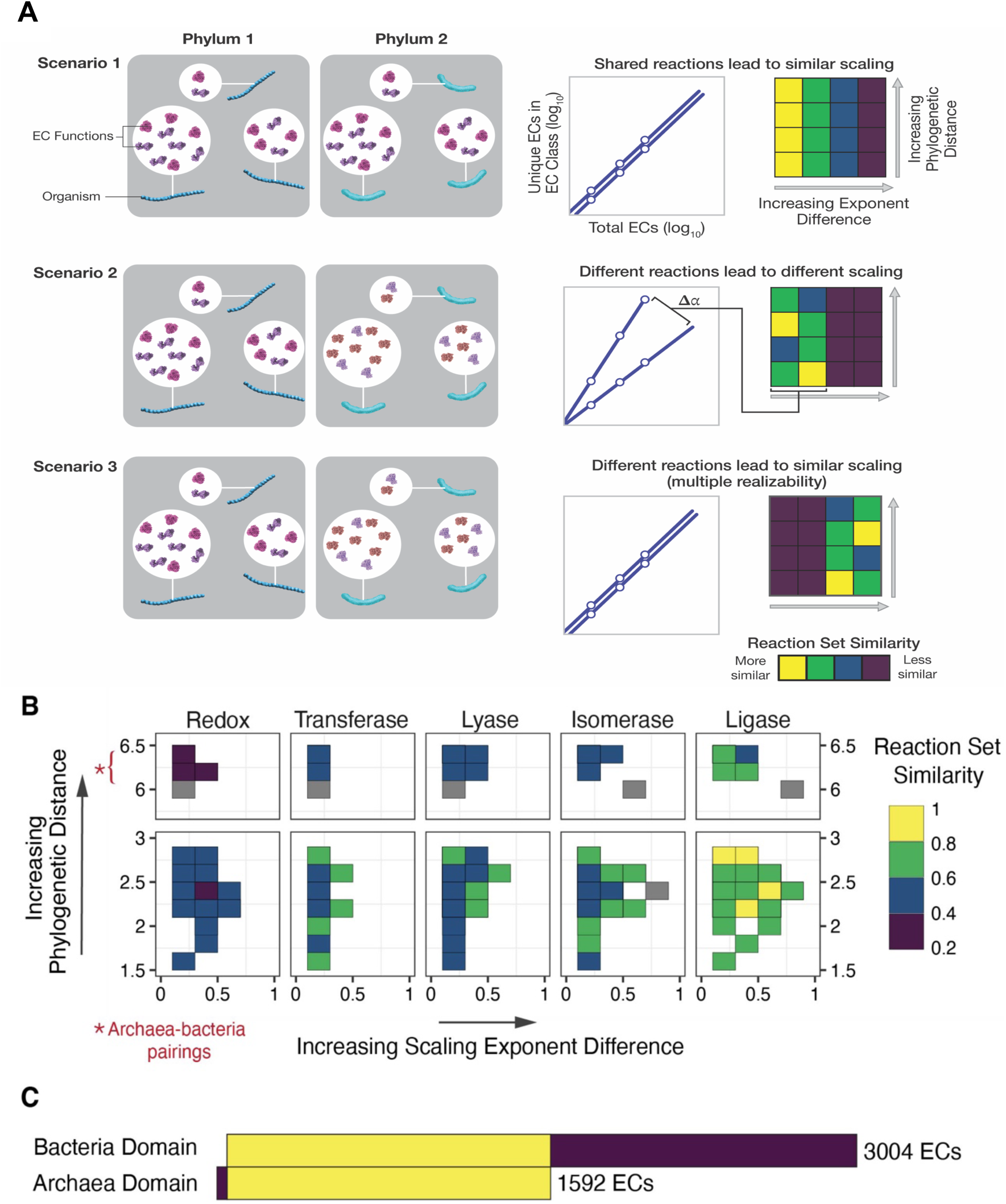
Similarity in reaction sets across bacterial and archaeal phyla do not explain convergence in functional scaling trends. **(A)** Schematic depicting different hypotheses for scaling trends. Scenarios 1 and 2 represent how similarity and differences in scaling could be explained by shared reactions among taxa. Scenario 3 describes an alternative scenario, where similar scaling arises through different reaction sets, providing evidence of multiple realizability. **(B)** Average reaction set similarities between phyla. Gray squares (NA) indicate the lack of size overlaps between the phylum pairs Deinococcota/Fibrobacterota and Deinococcota/Thermoproteota. **(C)** The overlap in four-digit EC codes in archaea and bacteria at the domain-level. Nearly all archaeal ECs are also found in bacteria.

Clustering phyla based on functional scaling could yield new hypotheses about the selective constraints acting on reaction diversity—constraints that we found are independent of ancestry or specific molecular substrates. We therefore compared different ways of clustering organisms beyond phylogenetics, which is history-based, to ones that are more convergence-based: (1) reaction set similarity and (2) functional scaling similarity. Reaction set similarity depicts phyla that are composed of similar reactions, which may reflect tight constraints that environmental or physiological context imposes on *specific* reactions that organisms can evolve. By contrast, similar functional scaling may also reflect tight constraints, but these are system-size dependent and reflect constraints on *ensembles* of reactions.

To determine similarity based on reaction sets, we constructed a dendrogram via a neighbor-joining algorithm, using average reaction set similarities as our distance matrix (see **Materials and Methods**). For functional scaling similarity, we constructed a separate dendrogram by hierarchically clustering phyla based on their five functional scaling exponents at the 1^st^ EC digit. We compared these two dendrograms to our adapted phylogenetic tree (introduced in **Figure 2C**). Surprisingly, we found no congruence in how phyla are grouped across the three methods (**Figure 4**). One might expect phylogenetic similarity to map to reaction similarity. However, we find this is not the case, corroborating earlier results supporting a decoupling between reaction similarity and evolutionary proximity^31^. Additionally, there are little to no preserved relationships among phyla even when considering functional scaling. A prominent example is the position of archaeal phylum Thermoproteota in each dendrogram: an outgroup in the phylogenetic tree, Thermoproteota is clustered with bacterial phyla in the other dendrograms, most closely with Chloroflexota in terms of reaction similarity, and with Pseudomonadota and Bacillota in terms of functional similarity. Overall, we propose new relationships between groups of organisms that capture common functional constraints on their system-size dependent reaction diversity, which differ from phylogenetic similarity or reaction similarity.

**Figure 4:**
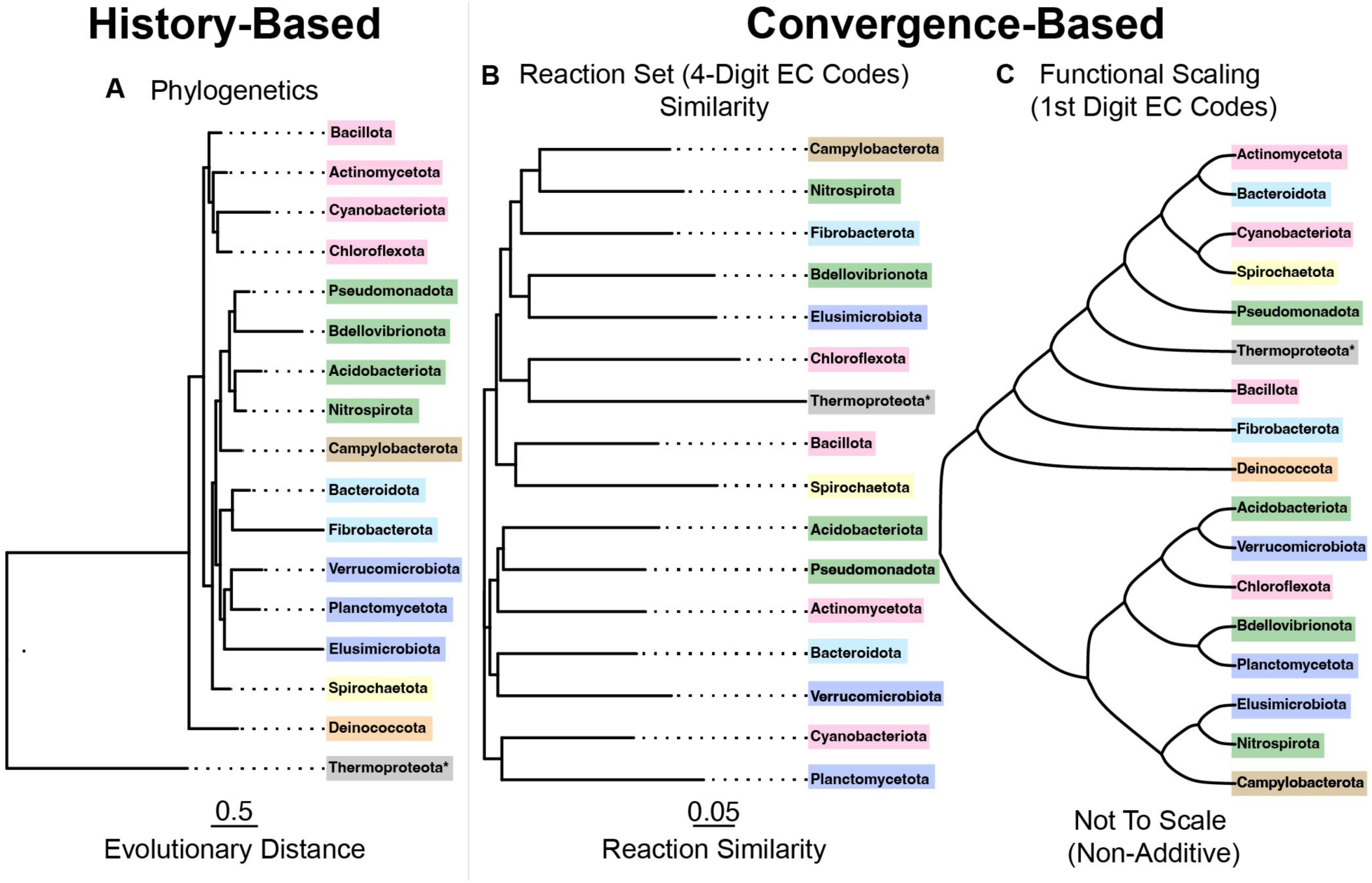
Dendrograms based on phylogenetics, four-digit EC codes, and functional scaling provide evidence for decoupling among ancestry, sets of reactions, and constraints in reaction diversity. **(A)** Phylogenetic tree based on similarity in conserved protein sequences, adapted from Martinez-Gutierrez and Aylward. **(B)** Neighbor-joining tree based on average reaction set similarity in taxa between phyla. Deinococcota is not included in this tree due to no size overlaps between this group and Thermoproteota/Fibrobacterota. **(C)** Hierarchically clustered dendrogram based on scaling exponents across **EC 1**, **EC 2**, **EC 3/4**, **EC 5**, and **EC 6**. The lack of congruence in how phyla are clustered across all three dendrograms reveals potentially novel relationships based on functional constraints in reaction ensembles.

### Physiology and Functional Scaling

A step in developing the use of functional scaling trends for predicting evolutionary outcomes will be identifying what environmental or physiological factors drive them. These factors become more accessible for finer-grained functional equivalence classes (2^nd^ and 3^rd^ EC digits), because these classes may vary more significantly across biological groups and some finer-grained classes may contribute more significantly to 1^st^ digit EC scaling behaviors than others (**SI Figure S4**). For this reason, we calculated scaling parameters for over 40 specific functional classes across bacteria, archaea, and eukarya (**Appendix, SI Data Files S3-S5)** at the 2^nd^ and 3^rd^ digit levels. We find scaling behaviors persist, but not in every class at every level of specificity (**Figure 5A**). Indeed, we find many differences across 2^nd^ and 3^rd^ digit functional scaling, suggesting that constraints on diversity do not operate equally on all types of reactions.

**Figure 5:**
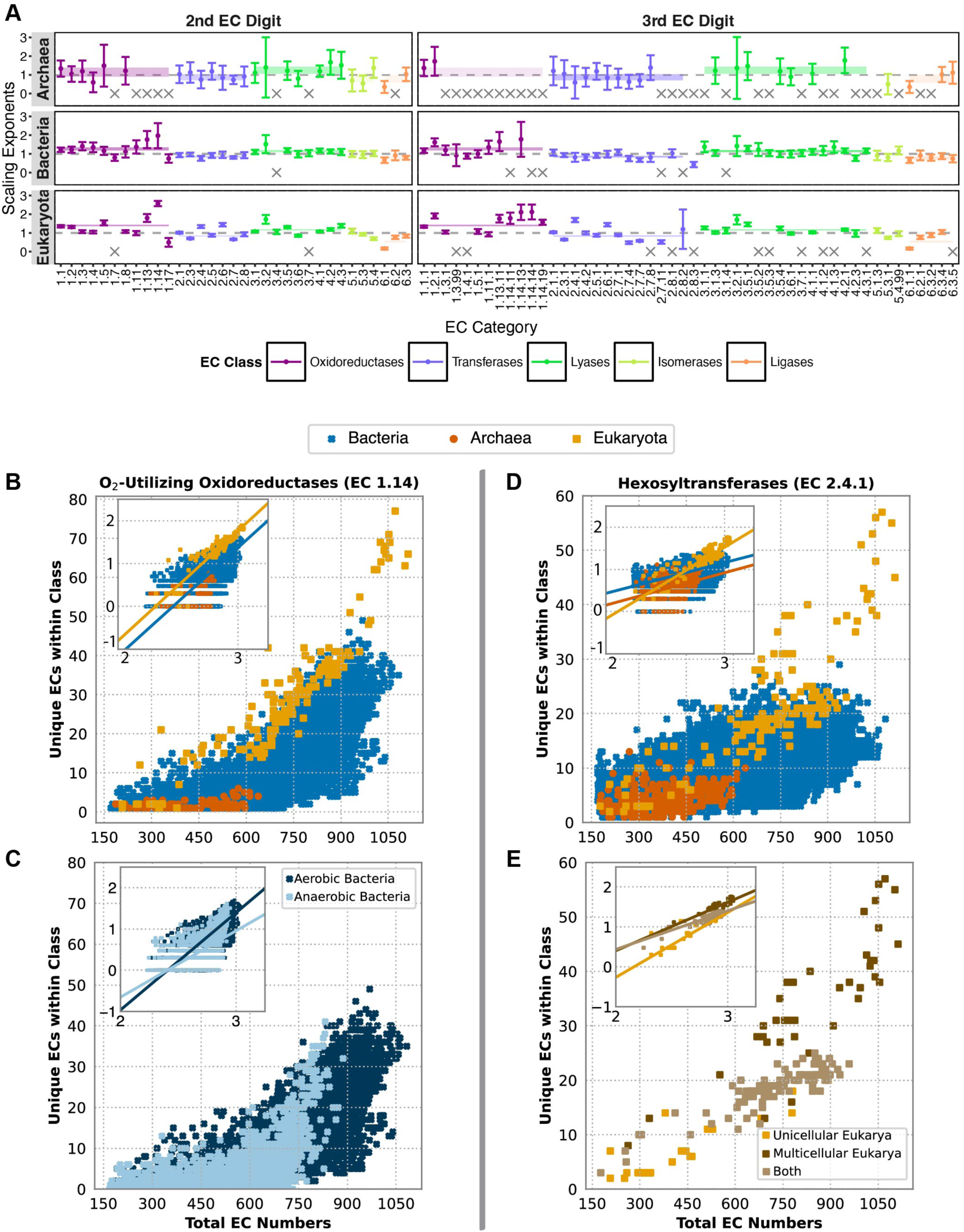
Scaling in specific functional equivalence classes indicate how at cell physiologies may contribute to constraining reaction diversity. **(A)** Scaling exponents across 2^nd^ and 3^rd^ digit functional equivalence classes suggest varying constraints on different biochemical functions. **(B)** Scaling of O_2_-utilizing oxidoreductases (**EC 1.14**), including log_10_-log_10_ transformed data (inset), varies across biological domains. (**C**) Mapping aerobic metabolism of bacteria to scaling trends in O_2_-utilizing oxidoreductases reveals a threshold for the total reactions encoded in anaerobic bacteria. (**D)** Scaling of hexosyltransferases (**EC 2.4.1**), including log_10_-log_10_ transformed data (inset), shows different scaling trends across domains (**E**) Mapping multicellularity of eukarya to scaling trends in hexosyltransferases demonstrates that multicellularity explains higher absolute diversity.

As a case study in mapping scaling trends to physiological factors, we chose two examples that exhibit their own distinct scaling behavior (or lack thereof) across domains. The first is a scaling trend emerging in a 2^nd^ digit class, the O_2_-utilizing oxidoreductases of **EC 1.14.-**. Enzymes in this class act on paired donors with incorporation or reduction of molecular oxygen, with 613 identified reactions. The second example is a 3^rd^ digit class, the hexosyltransferases (**EC 2.4.1.-**). Enzymes in this class transfer six-membered sugars like glucose from one molecule to another, with 364 identified reactions. Both functional classes significantly impacted the evolution of biochemistry: the evolution of biological O_2_-utilization added more energy for biochemical reactions^32^ and led to higher complexity and molecular diversity in biochemical networks^33,34^, while glycosyl-transferring reactions (**EC 2.4.-**), which encompass hexosyltransferases, are primarily responsible for the biosynthesis and breakdown of biomass on Earth^35^.

We hypothesized that the physiological constraints that enable the diversity of O_2_-utilizing reactions are associated with aerobic metabolism. **EC 1.14** reactions in bacteria and eukarya, which both contain aerobic taxa, scale superlinearly (2.508 ± 0.044, R^2^ = 0.531 for bacteria; 2.570 ± 0.140, R^2^ = 0.871 for eukarya), with eukarya exhibiting tighter variance (**Figure 5B**). Archaea show no scaling, in this case due to low absolute **EC 1.14** reaction diversity. Testing whether scaling differences result from cell physiology (i.e., whether superlinear behavior results from aerobic metabolism), we predicted which taxa can perform aerobic metabolism by tracing the presence of the EC codes most indicative of aerobic phenotypes^36^, focusing on bacterial taxa because these can be anaerobic or aerobic (**see Materials and Methods**). Though not a definitive classifier, this method performed moderately well in revealing expected systematic trends (**SI Figure S5**). We find for bacteria, there is a size threshold in anaerobic reaction diversity, where anaerobic bacteria can only support sizes of 800-900 unique reactions in total across all classes (**Figure 5C**). Additionally, we observe different scaling behaviors for aerobic vs. anaerobic bacterial taxa. Aerobic taxa exhibit steeper superlinear behavior (2.359 ± 0.046, R^2^ = 0.537) than anaerobic (1.610 ± 0.119, R^2^ = 0.238), though the low R^2^ value (below 0.5) for anaerobic samples suggest that scaling trends here are not strong. The scaling in aerobic samples follows more closely with eukaryotic and bacterial taxa overall. Taken together, these findings indicate that the presence of aerobic metabolism may help in explaining shifts in scaling exponents across domains, while also predicting a threshold to the total reaction diversity that bacterial cells can encode, even without considering the energy that O_2_-utilizing reactions add to metabolism.

Scaling trends for hexosyltransferases also indicate physiological constraints. The hexosyltransferases of eukarya scale superlinearly (1.689 ± 0.813, R^2^ = 0.813), with tighter variance and higher absolute diversity than those of bacteria: while large eukarya can have nearly 60 different encoded hexosyltransferases, similarly sized bacteria plateau at approximately 30 (**Figure 5D**). The low R^2^ values of the scaling fits for bacteria (0.726 ± 0.019, R^2^ = 0.284) and archaea (0.766 ± 0.177, R^2^ = 0.165) suggest that this functional equivalence class does not exhibit scaling constraints in these domains. These differences may be due to varying constraints in cell to cell signaling that arise in unicellular versus multicellular organisms. Transferring sugars between molecules plays a pivotal role in modifying proteins and lipids, thereby changing membrane composition and aiding in communication between cells^37^, which may be more important for large, multicellular eukaryotic cells than for unicellular archaea and bacteria. To investigate this further, we labelled eukaryotic taxa as multicellular, unicellular, or both (e.g., some fungi have dimorphic life cycles) to see if this would explain superlinear behavior in eukarya. We found that this was not true with our data: hexosyltransferases in multicellular taxa and ambiguous cases scale linearly (1.212 ± 0.233, R^2^ = 0.756 for multicellular; 0.967 ± 0.077, R^2^ = 0.835 for ambiguous), while those in unicellular samples counterintuitively scale superlinearly (1.626 ± 0.356, R^2^ = 0.724). However, further studies are needed to confirm these trends with more samples. These results indicate that there are shifts in scaling due to multicellularity, where multicellularity may limit the diversity of these functions as compared to unicellular samples. Additionally, it may be that other physiological features can explain scaling, such as the presence of more membrane-bound organelles in eukaryotes. Interestingly, we found that multicellularity does explain the higher absolute diversity in hexosyltransferases compared to their unicellular (or ambiguous) eukaryotic and bacterial counterparts at system sizes of 700 total reactions and above (**Figure 5E**), thereby connecting biochemical reaction diversity with higher-level physiological functions such as cell signaling.

One possible explanation for the increased reaction diversity within a functional class is that a specific protein fold diversified to catalyze all reactions within that class. This would mean functional scaling is attributable directly to protein structural diversity. However, we find that the diversity of reactions in our case studies of O_2_-utilizing enzymes and hexosyltransferases are not limited to specific protein folds. Both functional equivalence class examples are catalyzed by multiple types of folds, as indicated by the various CATH numbers associated to the relevant EC codes (**SI Tables S2 and S3**), which describe structural and evolutionary similarity among protein domains. In fact, reaction convergence is generally common among non-homologous proteins^38^. Thus, the scaling laws reported here are also not dictated directly by fold evolution, consistent with our findings that these scaling relations are not due to phylogenetic homology. Taken together our results indicate functional scaling is the result of emergent, system-size dependent ensemble constraints on reaction diversity in evolving systems, making the identified convergence potentially generalizable to other types of catalysts beyond enzymes.

## DISCUSSION

We identified over 120 new scaling relationships and showed how these arise due to convergence in the reaction diversity of life. Organisms that are distantly related, and composed of different sets of reactions, can nonetheless converge on the same functional scaling in one or more classes. Our results therefore support the multiple realizability of reaction sets that can fulfill the same ensemble-level physiological constraints. This suggests a previously hidden a layer of biochemical optimization operating on ensembles of enzymes, which is not reducible to the protein structures that might catalyze them or specific innovations in reaction chemistry. Even if the trajectories of protein catalysts are unpredictable, we have shown that the possible space of reactions that catalysts might collectively discover are tightly constrained and depend on the total reaction diversity encoded.

We aimed to illustrate a path forward for adapting these scaling relations as a predictive lens into the evolution of function in enzyme ensembles. To this end, we demonstrated system size-dependent patterns in two fine-grained functional equivalence classes, the O_2_-utilizing oxidoreductases of **EC 1.14** and the hexosyltransferases of **EC 2.4.1**. We anticipate other functional classes at the 2^nd^ and 3^rd^ digits will likewise be found to be driven by the physiological or environmental constraints that shape evolving enzyme populations. Indeed, the identified functional constraints could also be applied to other biochemical systems, such as viruses or other mobile genetic elements, which experience different selection pressures and have so far been difficult to phylogenetically characterize due to their rapid macromolecular evolution^39,40^.

The identified functional scaling trends can also be applied to design principles for engineering synthetic life, as they seem to impose physiochemical limits on how biochemical modules can be adaptively combined. For instance, taking account of functional scaling could aid in metabolic engineering, placing constraints on the combinatorics of integrating metabolic pathways^41^ and perhaps providing explanations for cases where integration may fail^42^. Our results can also inform directed evolution approaches. Understanding the emergent, predictable properties of co-evolving enzymes would allow novel steering of experiments involving many enzymes optimized together^43–45^. Additionally, a growing body of work is focused on designing controllable enzymatic networks that can perform certain behaviors like sensing, motility, and communication^46^. These, too, could be informed by functional scaling in their design, providing new tools for active matter research^47^. Our methods therefore offer new ways to inform adaptive properties of reaction networks broadly.

We showed the identified functional scaling trends do not depend on the exact identity of protein catalysts, nor reaction substrates. This suggests these convergent constraints might inform universal organizational principles of evolving reaction networks, because similar constraints might apply to other sets of evolving catalysts and substrates. For example, researchers could apply these scaling laws to theorizing about the earliest forms of life on Earth predating LUCA or to potential life on other worlds. All living chemical systems likely require ligation to build biomass and construct increasingly complex functional molecules, raising the question of whether ligation will control diversity of other reaction types in alternative or alien biochemistries, as we found it does in terrestrial life. Could this be true even in an entirely different class of catalysts that are distinct from proteins? Because of these scaling trends are not substrate dependent, the trends and their corresponding constraints could provide new insights into reaction networks that preceded the last universal common ancestor (LUCA), even before enzymes evolved. For this reason, we expect the existence of functional scaling relationships and trade-offs across reaction classes to inform origin of life research in areas as diverse as theoretical chemical modeling, autocatalytic sets and their reaction diversity^48–50^, experimental studies of dynamic combinatorial libraries^51,52^, and systems chemistry approaches^53,54^.

Despite the historical contingencies that impact enzyme structures, and the radically different environmental contexts that biological lineages must adapt to, our findings demonstrate that there are predictable regularities in biochemical evolution. The convergence of functional scaling trends suggests that observed patterns in reaction diversity are not a consequence of LUCA. We conjecture they may instead arise through other factors, such as a convergent feature of the physiologies and environment(s) of Earth. While it is currently unknown what possible chemistries could evolve on other worlds, functional scaling provides a new tool that could prove useful for predicting reaction diversity in evolving reaction networks, with applications to both Earth-based life and potential life on other worlds.

## MATERIALS AND METHODS

### Acquiring Data from Joint Genome Institute

We obtained our bacterial, archaeal, and eukaryotic genomic dataset from the Department of Energy-Joint Genome Institute (DOE-JGI) Integrated Microbial Genomes and Microbiomes (IMG/M Database)^24,25^, acquired in September 2023 through a text-mining Python script. This data included genome statistics (ex. total number of base pairs, total number of genes, total number of protein-coding genes, etc.), metadata, and Enzyme Commission (EC) number lists for each sample. Archaeal and bacterial data samples came from isolates, single-amplified genomes, and metagenome-assembled genomes, whereas eukaryotic data came strictly from isolates. Bacteria and archaea were scraped from the “JGI” category, whereas eukarya samples were scraped from the “All” category in the IMG/M database. Prior to filtering data, our data consisted of 19,653 bacteria, 843 archaea, and 686 eukarya samples.

### Filtering Data

We sought to retain samples with the highest quality of JGI function annotations. To this end, we filtered out samples whose protein-coding genes with function prediction was less than 10% or more than 90%, as even the most well-studied organisms do not have 100% functional annotation. Additionally, any gene without a protein-coding assignment that the JGI annotation pipeline mistakenly assigned an EC function was removed from the dataset. We excluded samples explicitly labeled “Fungal Annotation,” based on guidance from the Joint Genome Institute staff regarding potential reliability issues with this annotation category. However, we retained fungal samples annotated through other methods. We removed samples that fell under a determined threshold for number of genes, following the same thresholds as Gagler et al. 2022^15^. For archaea and bacteria, that organism was *Pelagibacter ubique,* which is the smallest free-living prokaryote discovered at the time of their analysis and consists of 1,354 genes^55^. For eukaryotes, the representative smallest free-living species was *Ashbya gossypii*, with 4,718 genes^56^. We removed any genomes whose total enzyme count was below 169 ECs, as this number represents the total EC count in the LUCA consensus model^57^ and serves as our proxy for minimal, non-parasitic life. Finally, we filtered out duplicate species that had the same distributions of ECs in order to reduce sampling bias. In each list of ECs, we removed partial ECs (‘1.1.1.-’) and ECs associated with glycans. Our final dataset post-filtering was 13,777 bacteria, 371 archaea, 203 eukarya.

### Determining Scaling Exponents from Empirical Data and Correlating with Phylogenetics

To calculate scaling parameters, we log-transformed our data using numpy (version 1.26.2) and generated an ordinary least squares (OLS) model using statsmodels (version 0.14.1). For the 1^st^ EC digits, we combined ECs 3 (hydrolases) and 4 (lyases) (see **Main Text** for details). In certain analyses, we prioritized equal representation of different phyla in domain-level scaling to reduce sampling bias inherent in biological databases. For bacteria and archaea, we examined phyla that have at least 40 taxa in them and that are represented in the phylogenetic tree created by Martinez-Gutierrez and Aylward, 2022^29^. We randomly sampled 40 taxa from each phylum and performed a scaling analysis on the resulting 760 bacterial datapoints and 80 archaeal data points, respectively. We performed this randomization 10 times and found the weighted average slope and confidence intervals. For archaea, the averaged slope of randomized trials and the slope when all filtered samples are considered fall within the confidence interval of the randomized version. For bacteria, this was true for ECs 1 through 5, but not EC 6 (**SI Figure S6**). Eukarya were not sampled in this manner, as only one phylum has at least 40 taxa (Ascomycota). Therefore, we retained all eukaryotic taxa to have a sufficient number of samples to perform scaling analyses on.

To find how scaling trends relate to evolutionary relationships, we investigated the correlation between scaling exponents and phylogenetic distances. Using the Python package DendroPy (v. 5.0.1) and a non-ultrametric tree representation from Martinez-Gutierrez and Aylward^29^, we calculated the phylogenetic distance between phyla by calculating the distance between every inter-phylum species pair and calculating the average. We conducted Spearman correlation tests using R (v. 4.4.0).

### Comparing Reaction Set Similarities Across Taxa

To compare reaction set similarities, we represented each taxon as a set of unique ECs, either for a given 1^st^ digit EC class in the case of **Figure 3B** or for all ECs total in the case of generating a dendrogram in **Figure 4B**. When analyzing only EC 1 oxidoreductase similarity, for example, a genome was represented as a set of unique ECs {1.1.1.1, 1.1.1.2, 1.1.1.4, … etc.}. To compare reaction set similarity at the phylum-level, we performed pairwise comparisons between taxa belonging to different phyla, deliberately excluding intra-phylum comparisons. We controlled for set size effects by restricting our analysis to cross-phylum pairs where taxa differed by no more than 10 total ECs. For each qualifying cross-phylum pair, we measured similarity by calculating their overlap coefficient, an adjusted Jaccard index that further controls for differences in set size.^58^ An overlap coefficient of ‘1’ indicates that reaction sets are either identical or that one set is ‘contained’ within another, whereas an overlap coefficient of ‘0’ indicates no shared reactions (four-digit EC codes) between the two sets. Finally, we calculated the average of cross-phylum comparisons to find an overall reaction set similarity at the phylum-level. Because this method depends on taxa having similar sizes (e.g., total number of encoded ECs), some phylum pair similarities could not be calculated, due to a lack of size overlaps (see **SI Figure S7**). These pairs were Deinococcota/Fibrobacterota and Deinococcota/Thermoproteota. See **SI Figures S8, S9, S10** for set comparisons among individual taxa, as well as EC appearances as a function of size.

### Constructing Phylogenetic, Reaction Set, and Functional Scaling-Based Dendrograms

To construct the phylogenetic tree at the phylum-level, we adapted a tree created by Martinez-Gutierrez and Aylward, which is based on concatenated ribosomal protein and RNA polymerase alignments with fine-grained phylogeny (multiple representative species per genus)^29^. Using R (v. 4.4.0) and libraries ggtree (v. 3.12.0) and ape (v. 5.8), we pruned the tree to only include phyla that are represented in our dataset and collapsed the tree to cluster species based on phylum^59^.

We constructed a dendrogram based on reaction set similarity through a neighbor-joining algorithm using Biopython (v. 1.84), inputting all pairwise overlap coefficients between phyla, where all taxa were represented as a set of *all EC codes* as the distance matrix. We removed ‘Deinococcota’ from this tree due to the lack of pairwise comparisons with Thermoproteota and Fibrobacterota, as taxa between these groups had no overlap in size (**see “Comparing Reaction Set Similarities Across Taxa”).**

We created a dendrogram that groups together phyla based on similarity in functional scaling behaviors at the 1^st^ digit EC level by performing a hierarchical clustering analysis with the Python package SciPy (version 1.11.4), where phyla were represented as vectors of the scaling exponent for the 1^st^ EC digit classes. We used Euclidean distances as the distance metric and Ward’s linkage method to prioritize reducing the internal variation in each cluster, adding confidence that the phyla that are clustered together have similar constraints driving their functional scaling behaviors. We found that dendrograms based on other linkage methods resulted in similar clusters despite differences in clustering statistics (**SI Figure S11, SI Table S4)**.

### Correlating Differences in Scaling Trends with Physiology

To understand the relationship between aerobic metabolism and scaling trends in O_2_-utilizing oxidoreductases, we labeled which taxa are likely aerobic or anaerobic by employing a method developed by Jabłońska and Tawfik in predicting aerobic phenotypes using only ECs^36^. They identified 5 ECs that were best at predicting aerobic phenotypes compared to other oxygen-utilizing enzymes. They are coproporphyrinogen oxidase (EC 1.3.3.3), nitric-oxide synthase (EC 1.14.14.47), stearoyl-CoA 9-desaturase (EC 1.14.19.1), FAD-dependent urate hydroxylase (EC 1.14.13.113), and aklavinone 12-hydroxylase (EC 1.14.13.180), the latter of which was not represented in our JGI data. We designated a taxon as likely aerobic if it possesses at least one of these ECs and likely anaerobic if they did not.

For the relationship between multicellularity and hexosyltransferases, we labeled eukaryotic taxa as unicellular, multicellular, or both (e.g., dimorphic lifestyles) by examining what types of species each phyla contained (**SI Figure S12**). For instance, all taxa in the Chordata phylum were labeled multicellular, as this phylum only contains animal species. Taxa in a phylum like Ascomycota, relating to yeast species, were designated “both” for changes in multicellularity in their life cycle. These physiological assignments can be found in **SI Data File S6**.

## ACKNOWLEGEMENTS

We graciously thank Alexander Sonke, L. Felipe Benites, Riddhi Gondhalekar, Elizabeth Trembath-Reichert, Jordan Okie, Dániel Czégel, Hyunju Kim, and the Emergence Lab at Arizona State University for insightful discussions, critical feedback, and helpful guidance on biological and statistical analyses for this project. We thank artist Mesa Schumacher for refining our schematic visualizations for Figures 1 and Figure 3A. We are grateful to Carolina Martínez-Gutiérrez for providing phylogenetic data from previously published work and guidance on constructing our adapted tree. The authors kindly thank ASU Research Computing for HPC support. Portions of the data analysis code were generated with the assistance of GitHub CoPilot and ChatGPT. This work was supported by NASA Interdisciplinary Consortia for Astrobiology Research (ICAR) grant number 80NSSC21K1402.

## AUTHOR CONTRIBUTIONS

The initial conceptualization, supervision, and funding of this work was provided by S.I.W. V.M., C.M., and C.P.K. contributed to conceptualization. V.M. performed all data analyses and developed all code. V.M., S.I.W. wrote the manuscript with input from C.P.K. and C.M.

